# The meaning of the Michaelis-Menten constant: K_m_ describes a steady-state

**DOI:** 10.1101/608232

**Authors:** Enric I. Canela, Gemma Navarro, José Luis Beltrán, Rafael Franco

## Abstract

Often, in vitro or in vivo enzyme-mediated catalytic events occur far from equilibrium and, therefore, substrate affinity measured as the inverse of ES ⇄ E+S dissociation equilibrium constant (*K_d_*) has a doubtful physiological meaning; in practice it is almost impossible to determine *K_d_* (except using stopped-flow or other sophisticated methodologies). The Michaelis-Menten constant (*K_m_*), the concentration of substrate ([S]) providing half of enzyme maximal activity, is not the (*K_d_*). In the simple E+S ⇄ ES → E+P or in more complex models describing S conversion into P, *K_m_* must be considered the constant defining the steady state at any substrate concentration. Enzyme kinetics is based on initial rate determination, i.e. in the linear part of the S to P conversion when the concentration of [ES] remains constant while steady state occurs. We also show that Systems Biology issues such as the time required to respond to a system perturbation, is more dependent on *k*_1_, the kinetic constant defining substrate-enzyme association, than on *K_m_*. Whereas *K_m_* is instrumental for biochemical basic and applied approaches, in any physiological condition, an important parameter to be considered is the substrate association rate (*k_1_*).

## Introduction

At Chemistry, Biology, Pharmacy and even Medicine schools, enzyme kinetics is taught according the visionary work of Briggs and Haldane and of Michaelis and Menten [1]. The key formula given to students consists of a hyperbolic relationship between enzyme activity (v) and substrate concentration ([S]):

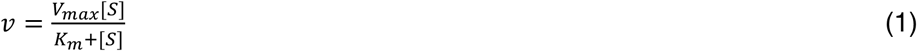

where V_max_ is the maximal activity and *K_m_* the Michaelis constant. *K_m_* is a function of the kinetic constants of the elementary steps in:

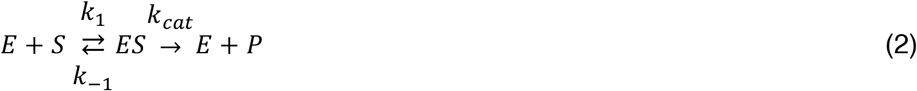

being *k_(i)_* the kinetic constant of every step, *K_m_* becomes:

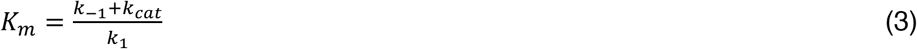

At the beginning of the 20th century Michaelis-Menten [2] did pioneer enzymology research and provided tools to calculate related parameters. On the one hand, the report was written in German and, accordingly, only available to those able to read in this idiom. Fortunately, the work of Johnson and Goody [3] included the translation of the original work, thus being relevant for scientists to make them aware of the message provided by Michaelis and Menten. On the other hand, the way of presenting data was very different to that used 100 years later.

Therefore, many enzymologists may not be aware of the meaning of the parameter “invented” by Michaelis and Menten, which was not *K_m_*. As described by Johnson and Goody [3]: “*Rather, they derived V_max_*/*K_m_, a term we now describe as the specificity constant, k_cat_/K_m_, multiplied by the enzyme concentration*…”.

For the purpose of this article it does not matter the actual concentration of the enzyme, but to simplify, we use [E] = 1. Then, the specificity constant, *k_cat_*/*K_m_*, provided by Michaelis and Menten [2] would be: *k_cat_*/*K_m_* ·1= *k_cat_*/*K_m_*.

## Results and Discussion

### Revisiting procedures for *K_m_* calculation: The Michaelis-Menten paradox

One paradox within the so-called Michaelis-Menten approach consists of deciphering *K_m_*’s mechanistic meaning; the challenge being to solve it for validity in *in vitro* and *in vivo* conditions. *K_m_* is calculated in isolated systems measuring initial rates at different substrate concentrations and fitting data to the Michaelis-Menten equation.

Back for decades, linearizations (e.g. the Lineweaver-Burk linearization [4], reported 20 years after the Michaelis-Menten paper) were instrumental for parameter determination. As commented elsewhere [5], the Eadie-Hofstee linearization introduces much less error in parameter estimation. At present, data must be fitted directly to the Michaelis-Menten equation using non-linear regression. But which are the data to be fitted? In brief, the reaction reate (v) for each substrate concentration must be taken from initial values, i.e. using the slope of the linear part of the plot of substrate disappearance (or product formation) versus time. Thus, data consists of pairs of substrate concentration and slopes of increment of product (or decrement of substrate) with time.

#### Interplay between *K_m_, k_1_, k_−1_* and *k_cat_* values

Our first aim was to understand the range of *k_1_* values when *k_cat_* is higher, similar or negligible in comparison with *k_−1_*. We have used the equation 1 to calculate several v/[S] data pairs for *K_m_* = 50 μM and *k_cat_* = 300 s^−1^. Considering equation 3, we calculated k_1_ values using the following parameters: i) *k_cat_* = *k_−1_*/10, ii) *k_cat_* = *k_−1_*/5, iii) *k_cat_* = *k_−1_*/2, iv) *k_cat_* = *k_−1_*, v) *k_cat_* = 2·*k_−1_*. The calculated k_1_ values are shown in table 1.

**Table 1.** k_1_ and K_d_ values calculated for K_m_ = 50 μM and k_cat_ = 300 s^−1^, and varying k_−1_ from 3000 to 150 s^−1^

| $k_{-1}(\text{s}^{-1})$ | 3000 | 1500 | 600 | 300 | 150 |
| --- | --- | --- | --- | --- | --- |
| $k_1(\text{M}^{-1}\text{s}^{-1})$ | $6.67 \cdot 10^7$ | $3.66 \cdot 10^7$ | $1.87 \cdot 10^7$ | $1.27 \cdot 10^7$ | $9.00 \cdot 10^6$ |
| $K_d$ | $4.50 \cdot 10^5$ | $4.10 \cdot 10^5$ | $3.21 \cdot 10^5$ | $2.36 \cdot 10^5$ | $1.67 \cdot 10^5$ |

If *K_m_* is fixed at 25 μM and *k_cat_* is kept at 300 s^−1^, the trend is similar and the respective *k_1_* values in the 3000–150 range of *k*_−1_ values (as in table 1) are: 1.3·10^8^, 7.2·10^7^, 3.6·10^7^, 2.4·10^7^ and 1.8·10^7^ M^−1^s^−1^. These results indicate that *k_cat_* going from similar or significantly lower (1/10) values than *k_−1_* do not severely impact on *k_1_* values. In fact, at either 25 or 50 μM *K_m_* value, a 20-fold change in *k_cat_*/*k_−1_* ratio results in a change of only 7-fold in *k_1_* values.

In general terms, and using the simplest mechanism, *K_d_* is lower than *K_m_*. Fig. 1 shows that *K_d_* is much lower thant *K_m_* when *k_cat_* is 100 times greater tan k_−1_ (*k_cat_*/*k*_−1_=100) and that it is required a change of three orders of magnitude (*k_cat_*/*k_−1_*= 0.1) for convergence of *K_m_* to *K_d_* values.

**Fig. 1.**
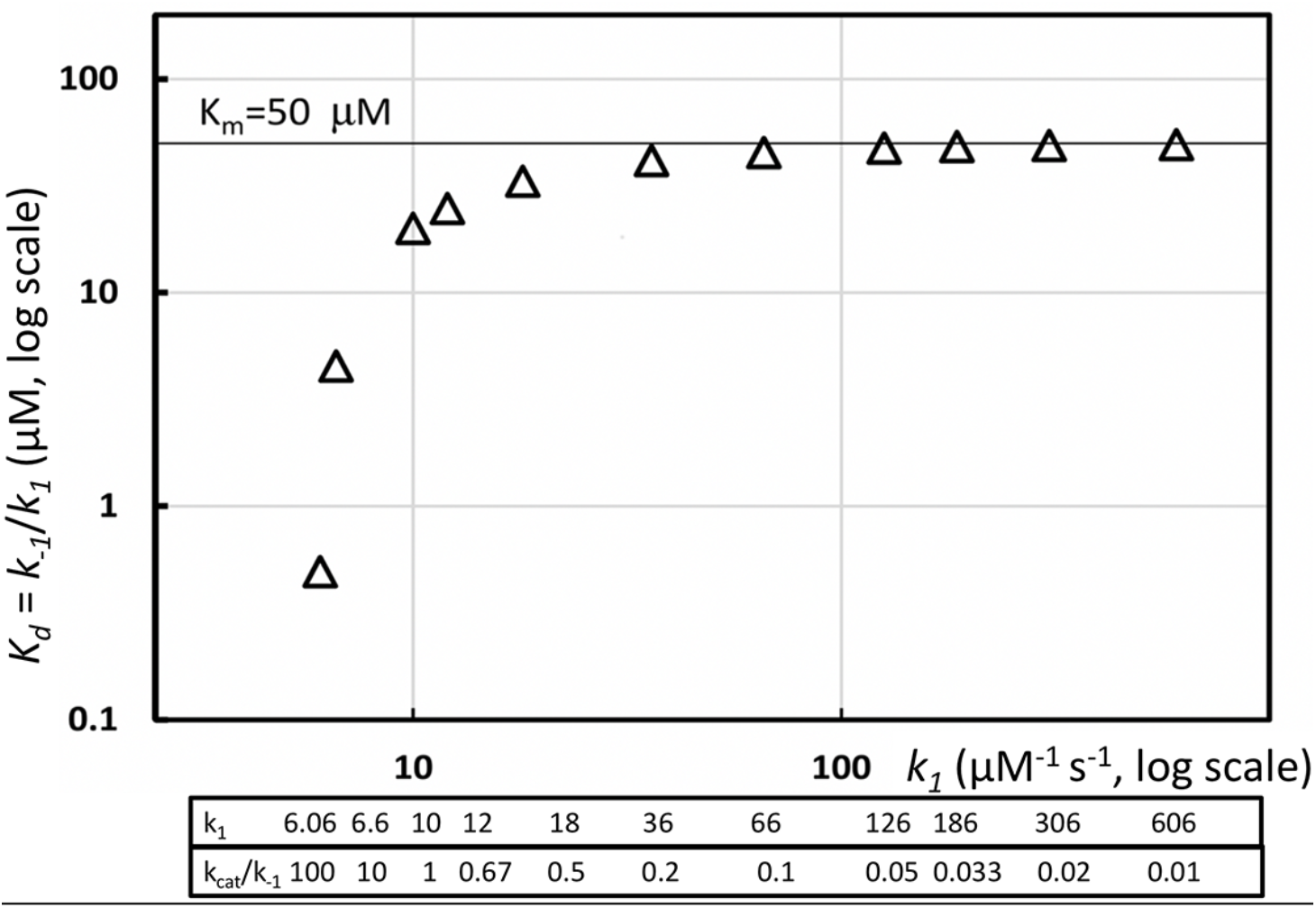
Dissociation equilibrium constant for different values of k_1_ values at different at different k_cat_/k_−1_ ratios (bottom). k_cat_ and were fixed at *K_m_*, respectively, 300 s^−1^ and = 50 μM. Data points were obtained using eq. 3.

In summary, *K_m_* being close to an equilibrium constant requires low values of *k_cat_* with regard to *k_−1_*. Do they really occur in either *in vitro* or *in vivo* conditions? *k_cat_* is the number of molecules of product formed per one molecule of enzyme and time unit. Enzymes are very efficient and *k_cat_* values may be measured in hundreds/thousands per second. Obviously, exceptions occur and k_cat_ may be lower (e.g., structurally complex substrates). We had extensive experience with adenosine deaminase, whose congenital deficit produces severe combined immunodeficiency [6]. In this case, *k*_*ca*t_ is in the order of hundreds in s^−1^ units (251), for the most common substrate, adenosine, and 283 for a structural analog, 2’-deoxiadenosine [7,8]. *K_m_* values for these compounds are in the 20-30 μM range, in close agreement with our own results [9,10]. 250 molecules of adenosine transformed every second by one enzyme molecule is not negligible in absolute terms. May 250 s^−1^ be negligible due to the fact that the k_−1_ value is significantly higher? Using stopped-flow spectrofluorometer measurements using pure adenosine deaminase from calf intestine, it was reported [7] that *k_cat_* = 244 s^−1^, *k_1_* = 3.1·10^7^M^−1^s^−1^ and *k_−1_* = 500 s^−1^. Then *k_cat_* is not negligible in front of k_−1_, and the constants would be *K_d_* = 16 μM and *K_m_* = 24 μM, i.e. *K_m_* would be a 33% higher than *K_d_*. While the k_cat_ value is easy to calculate, technical issues make difficult to make reliable estimations of *k_1_* and *k_−1_*, and any miscalculation in these rate constants would affect the distance between *K_m_* and *K_d_* values. Hence, the message is that two substrates having similar *K_d_* values, the one with lower *k_1_* will lead to a higher *K_m_*, and vice versa. If should be noted that in complex mechanistic scenarios *K_m_* may be (at the theoretical level) lower than *K_d_* (see below).

### Fitting enzyme kinetic data by non-linear regression using the classical expression for *K_m_*

Using the classical model: E+S=ES➔E+P, the substrate with quickest association kinetics will display a lower *K_m_* in *in vitro* assays. Our next aim was to fit in silico-generated data pairs of v/[S] to obtain *K_m_* by non-linear regression fit to the Michaelis-Menten equation, or to the equation obtained from substituting *K_m_* by the expression indicated in equation 3:

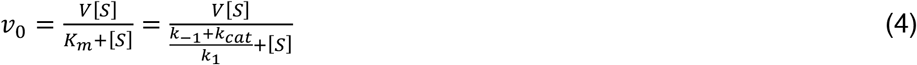

In both cases the parameters were refined by using the macro ref_GN_LM [11], which consists of an adaptation of Levenberg-Marquardt modification of the Gauss-Newton iterative algorithm [12], for use under the MS Excel™ spreadsheet. In order to compare results, we have tested two different objective functions, the first one defined as the sum of squared errors (U); in the second case, the function to be minimized is the sum of squared relative errors (U_rel_):

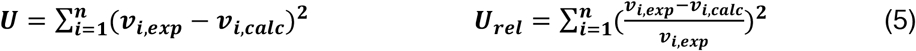

where *n* is the number of data points, *v_i,exp_* the experimental value of the *i*-point, and *v_i,calc_* the calculated after the model (eq. 4, center or eq. 4, right).

#### Data fitting

When fitting was performed using eq. 4, center, which has 2 parameters, we obtained the values used to generate the data (*K_m_* = 50 μM and *k_cat_* = 300 s^−1^). Results were almost identical using one of the other minimization procedure (*K_m_* = 50, SD = 0.01 and *k_cat_* = 300, SD = 0.01 for U, and *K_m_* = 50, SD = 0.03 and *k_cat_* = 300, SD = 0.14 for U_rel_). When 3 parameters were considered (eq. 4, right), fitting results were inconsistent (see table 2). Even when *k_cat_* was fixed to 300 s^−1^ the data still were not consistent, as *k*_1_ is 6 but *k*_−1_ becomes close to 0 and with huge SD values. The parameters and the SDs, calculated under different initial contour conditions are given in table 2. In summary, the robust fitting of data to eq. 4, center, shifts to a non-reliable fitting when using eq. 4, right, thus suggesting both that data do not allow calculation of *k_1_* and *k_−1_* and that *K_m_* cannot be an equilibrium constant. This fitting exercise using eq. eq. 4, right, in fact has not any rigor, as a “Michaelian” v/[S] plot leads to an equilateral hyperbola that is defined by just 2 parameters; hence, fitting to 3 (or more) parameters is impossible or, in other words, a vain actuation: the equilateral hyperbola cannot be described by 3 parameters (*k_1_, k_−1_* and *k_cat_*) but by 2. Leaving aside any “Michaelis-Menten paradox” it is necessary to continue to work using the canonical model and the current approaches albeit with caution. On the one hand, *K_m_* cannot be any more considered as a measure of substrate affinity. It should be noted that if *K_d_* is the equilibrium dissociation constant of the ES complex in the model described in eq. 2, and assuming steady state conditions, one can analytically devise eq. 3, which shows that only if *k_cat_* ≪ *k_−1_, K_m_* = *k_−1_*/*k_1_* = *K_d_*. The model predicts that *K_m_* can be relatively close to *K_d_* if *k_cat_* < 0.05·*k_−1_* (see Figure 1). This fact is known [5], but it may be overlooked: *k_cat_* is non-negligible for the majority of enzymes.

**Table 2.** Parameters after fitting data to eq. 4, right, and using the data generated with K_m_ = 50 μM and V = 300, [E] = 1 nM (Relative Units). The estimated standard deviation (SD) is given in parentheses.

| Initial values | Error type* | $k_{\text{cat}}$<br>($\text{s}^{-1}$) | $k_1$<br>($\mu\text{M}^{-1}\text{s}^{-1}$) | $k_{-1}$<br>($\text{s}^{-1}$) |
| --- | --- | --- | --- | --- |
| $k_1 = 0.01$<br>$k_{-1} = 10^{-5}$<br>$k_{\text{cat}} = 1$ | A | 300 (0.01) | 6.0 (7E-4) | 5.4E-9 (6.7E-6) |
| $k_1 = 0.01$<br>$k_{-1} = 10^{-5}$<br>$k_{\text{cat}} = 1$ | R | 300 (0.15) | 6.0 (0.002) | 5.8E-9 (4.9 E-5) |
| $k_1 = 100$<br>$k_{-1} = 100$<br>$k_{\text{cat}} = 100$ | A | 300 (0.015) | 26.1 (1.0) | 1010 (50.2) |
| $k_1 = 100$<br>$k_{-1} = 100$<br>$k_{\text{cat}} = 100$ | R | 300 (0.21) | 22.4 (4.3) | 818 (216) |
| $k_1 = 1$<br>$k_{-1} = 1$<br>$k_{\text{cat}} \text{ fixed (300)}$ | A | - | 6 (0.5) | 0.099 (26) |
| $k_1 = 1$<br>$k_{-1} = 1$<br>$k_{\text{cat}} \text{ fixed (300)}$ | R | - | 6 (0.049) | 6E-6 (0.049) |

\* A: absolute error; R: relative error (eq.5)

Adding mechanistic complexity does not solve the paradox. For instance the introduction of a further step and consideration of conformational changes during catalysis (as described elsewhere [13,14]):

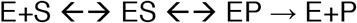

makes more complex the formula of *K_m_* while the calculation of the constant includes the fitting to an equation of only 2 parameters (equilateral hyperbola indicated in eq. 1). Then the message does not change except in that in these complex systems *K_m_* may be lower than *K_d_* if the las step EP → E+P occurs very slowly, i.e. with extremely low *k*_*ca*t_: something rather unusual.

### Is *K_d_* measurement feasible and has *K_m_* any mechanistically significant meaning?

The so-called Michaelis-Menten development in enzyme kinetics has been very useful despite enzymes do not work near any real equilibrium. Calculating the equilibrium constant of S ➔P reaction is simple: it is a matter of waiting for the equilibrium to occur and do the quotient [P]_eq_/[S]_eq_. Equilibrium dissociation constant for ES ⇄ E+S is a mechanistic resource but of limited value; it does not provide much information: when the enzyme is acting there is not any real equilibrium. In addition, it is very difficult to calculate with reliability. For this reason, *K_m_* has been often used as a substitute but we argue that this is not necessary and/or is of limited usefulness.

*K_m_* is a value obtained from v/[S] plots, and is equivalent to the concentration that provides 50% of V_max_ ([S]_0.5_). The classical mechanistic model (eq. 3) assumes that i) steady state occurs for each [S] and ii) [enzyme] is negligible (<1000·[S]). Taking these precautions, *in vitro* assays leads to a linear relationship between substrate consumption and time that, by definition, lasts until the steady state is no longer valid. At a given [S]: v = *k_cat_* [ES] and, therefore, v is constant only when [ES] is constant. From this fact two different approximations may be derived.

In terms of mechanisms, let us consider the two most common assumptions, for instance a rapid E + S = ES equilibrium; that is, the concentration of ES does not vary because *k_1_* and *k_−1_* are high and the equilibrium is achieved in few seconds. Then, both *k_1_* and *k_−1_* are higher than *k_cat_* and *K_m_* would be equivalent to *k_−1_*/*k_1_* (*K_d_*). As discussed below, *K_m_* cannot be an equilibrium constant as no equilibrium has been reached.

An alternative option is the so-called quasi steady-state approximation (QSSA), introduced in enzymology by Briggs and Haldane [15]. They assumed that, in the model defined in eq. 1, within the period of time that the [ES] remains constant: *k_−1_* ≪ *k_1_* and *k_−1_* is negligible in front of *k_cat_*. Whereas rapid equilibrium leads to a *K_m_* ≈ *K_d_*, QSSA leads to a *K_m_* ≈ *k_cat_*/*k_1_*, which has nothing in common with an equilibrium constant. Whereas in *vitro* assays are performed in conditions of negligible enzyme concentration ([enzyme] < 1000·[S]), enzyme and metabolite concentrations are not so distant in a physiological situation.

Surely, due to procedural factors, enzymologists have used *K_m_* values to compare “affinities” of different compounds using sentences such as “*the substrate with lower K_m_ has more affinity for the enzyme*”. This may happen as an exception for different reasons being Briggs and Haldane’s QSSA the most likely. In short, *K_m_* is indeed instrumental in Enzymology but cannot be used to define substrate affinity. The crucial question then is whether *K_m_* is used in a reliable manner when introduced in any Systems Biology study/analysis.

### Impact of the steady-state constant on Systems Biology approaches

The steady-state may be fulfilled in *in vitro* conditions, but may be a rare phenomenon in physiological *in vivo* situations [16–19]. Enzymes are parts of a system; accordingly, real *K_m_* value and meaning should be closely scrutinized in *in vivo* scenarios. In fact, each enzymatic step provides independent variables in Systems Biology approaches involving metabolic pathway-related calculations. We claim that the meaning of *K_m_* is apprehended in a “dynamic” framework. The conception may appear as trivial but conceptually what we propose is that *K_m_* is equivalent in a dynamic situation to *K_d_* in a static situation. Irrespective of the numeric values, *K_d_* is the dissociation constant (of the reaction E + S = ES) and *K_m_* is the “steady-state” constant. Thus, *K_d_* = [E]_eq_[S]_eq_/[ES]_eq_, and *K_m_* = [E]_ss_[S]_ss_/[ES]_ss_, “eq” standing for equilibrium and “ss” standing for steady state. If [E]_T_ is the total amount of enzyme, we may consider:

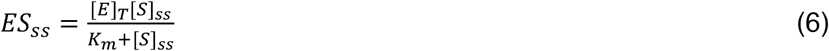

An expression that is valid for any substrate concentration used in *in vitro* assays. It is noteworthy that, for a given *k_cat_*, the greater the *K_m_*, the lower [ES]_ss_ and, consequently, the lower the reaction rate. In any physiological system the majority of reactions are far from equilibrium and the flux (e.g. glycolytic versus gluconeogenic or vice versa) goes in one direction. For such reactions *k_−1_* is negligible in front of *k_cat_*, and in consequence, for a given *k_cat_*, the higher the *k_1_* the lower the *K_m_* and the higher the flux provided by the catalytic step. In other words, in many *in vivo* physiological conditions, fluxes depend on *k*_1_.

In steady state conditions as those used in the pioneering work by Kacser and Burns [20–22] to study metabolic control, when the system is perturbed by adding a small amount of a substrate S, the parameter that measures the change in the flux, the reaction velocity, is known as elasticity. Taking an unbranched metabolic system and being E a member of the reaction chain, an increase in the input leading to a change in metabolic flux leads to two phenomena that are deduced by the above considerations. The value of the flux at the new steady-state will be directly proportional to [ES]. Obviously, this applies to all enzymes in the unbranched metabolism. Elasticity, as elsewhere defined [20,23,24] is given by:

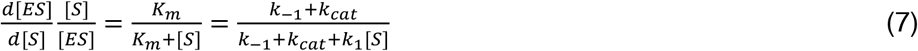

Therefore, the higher the *K_m_* value the lower the effect of a differential change of [S] on [ES]_ss_. Again, if *k_cat_* ≫ *k_−1_*, then elasticity becomes: *k_cat_*/(*k_cat_*+*k_1_*[S]_ss_). Thus, the greater the *k_1_*, the lesser the relative impact of [S] variation in the flux.

The second phenomenon concerns time, i.e. it is related to the kinetics of achieving of a new steady state when a perturbation to the system is applied, namely when the value of a metabolite (substrate) concentration changes. Being [E]_T_ the amount of enzyme, from an initial state (A): [ES]_A_ = [E]_T_[S]_A_/(K_m_ + [S]_A_), a perturbation (a change of the substrate concentration) will lead to a new state (B), where:

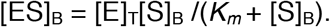

Taking the canonical model described in eq. 2), in which d[ES]/dt=k_1_[E][S]−(k_cat_+k_−1_)[ES], analytical integration is leads to the following relationship:

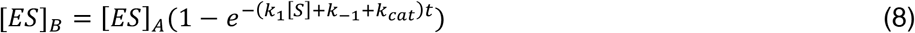

Thus, any perturbation in a step within a metabolic system will lead to a new steady state that depends on the [S] in the new steady state and on the kinetic constants of the enzymatic reaction. In eq. 8, *t* would be the time necessary to achieve the new steady state. Such variation in [ES] from steady state A to steady state B, if experimentally followed with reliability, will provide a specific constant (*k_obs_*) whose value is the exponent: *k_obs_* = *k_1_*[S]+*k_−1_*+*k_cat_*. If *k_−1_* is negligible versus *k_cat_* the equation becomes: *k_obs_* =*k_1_*[S]+*k_cat_*. Hence, *k_obs_* for a given metabolite concentration depends on *k_1_* of the corresponding enzyme. Elasticity gives information on the proportion of enzyme molecules that are occupied, and therefore engaged in the catalysis. The higher the *K_m_* value the lower the degree of occupation (lower the [ES]) and elasticity gives us an idea on how [ES] will change when [S] changes. In summary, elasticity measures the change while the kinetics to achieve the new steady state is defined by *k_obs_*. The take home message from this latter part of the article is that depending on the actual value to metabolite (substrate) concentration, the new steady state would depend i) on *k_1_*[S], ii) on *k_cat_* or iii) on *k_1_*[S]+*k_cat_*.

For instance, in cases of low [S], the kinetics in achieving a new steady state will depend on *k_cat_* (i.e. in an intrinsic property of the enzyme). Similarly, if the enzyme displays high *k*_*ca*t_ values, the kinetics from passing to a different steady state will be similar within a wide range of substrate concentrations. Data calculated without any a priori assumption and using the above values for *K_m_* and *k_cat_* (table 3), tells that both *k*_1_ and [S] affect *k*_obs_. At [S] = *K_m_* (50 μM), the time in sec needed to pass to 70 μM is 1.2·10^−5^ for *k_1_* = 6.67·10^7^ M^−1^s^−1^ and 8.88·10^−5^ for *k_1_* = 9.0·10^6^ M^−1^s^−1^. Interestingly, the time to reach a new steady state varies in a greater proportion than the [S]; when *k_1_* = 6.67·10^7^ M^−1^s^−1^ and initial [S] = 3.0·10^−5^ M, the time to reach 5.0·10^−5^ M is 4.34·10^−5^ s, whereas if the starting substrate is 5 times higher, ([S] = 1.5·10^−4^ M), the time to pass to 1.7·10^−4^ M is about two orders of magnitude higher (7.45·10^−7^ s).

**Table 3.** Determination of time needed to attain a new steady state. For a given [S] in the table the value of [S] at the simulated new steady state would be 20 μM higher than the “starting” steady state, which was defined by 10 μM [S] and 0.17 M [ES]_ss_ calculated using the same values used earlier for K_m_ (50 μM), k_cat_ (300 s^−1^) and E_T_ (1 nM). k_obs_ and time (t) to attain a new steady state were calculated (see text for details) using different k_1_ and k_−1_ values (note that fixing either value, the second one is related by the formula K_m_ =(k_−1_+k_cat_)/k_1_).

| [S]<br>(M) | [ES]<sub>ss</sub><br>(nM) | $k_1 = 6.67 \cdot 10^7$<br>($\text{M}^{-1}\text{s}^{-1}$) | | $k_1 = 3.67 \cdot 10^7$<br>($\text{M}^{-1}\text{s}^{-1}$) | | $k_1 = 1.87 \cdot 10^7$<br>($\text{M}^{-1}\text{s}^{-1}$) | | $k_1 = 1.27 \cdot 10^7$<br>($\text{M}^{-1}\text{s}^{-1}$) | | $k_1 = 9.00 \cdot 10^6$<br>($\text{M}^{-1}\text{s}^{-1}$) | |
| --- | --- | --- | --- | --- | --- | --- | --- | --- | --- | --- | --- |
| | | $k_{\text{obs}}$<br>($\text{s}^{-1}$) | t<br>(s) | $k_{\text{obs}}$<br>($\text{s}^{-1}$) | t<br>(s) | $k_{\text{obs}}$<br>($\text{s}^{-1}$) | t<br>(s) | $k_{\text{obs}}$<br>($\text{s}^{-1}$) | t<br>(s) | $k_{\text{obs}}$<br>($\text{s}^{-1}$) | t<br>(s) |
| $3.0 \cdot 10^{-5}$ | $3.75 \cdot 10^{-1}$ | $5.33 \cdot 10^3$ | $4.34 \cdot 10^{-5}$ | $2.93 \cdot 10^3$ | $7.90 \cdot 10^{-5}$ | $1.49 \cdot 10^3$ | $1.55 \cdot 10^{-4}$ | $1.01 \cdot 10^3$ | $2.29 \cdot 10^{-4}$ | $7.20 \cdot 10^2$ | $3.22 \cdot 10^{-4}$ |
| $5.0 \cdot 10^{-5}$ | $5.00 \cdot 10^{-1}$ | $6.67 \cdot 10^3$ | $1.20 \cdot 10^{-5}$ | $3.67 \cdot 10^3$ | $2.18 \cdot 10^{-5}$ | $1.87 \cdot 10^3$ | $4.28 \cdot 10^{-5}$ | $1.27 \cdot 10^3$ | $6.31 \cdot 10^{-5}$ | $9.00 \cdot 10^2$ | $8.88 \cdot 10^{-5}$ |
| $7.0 \cdot 10^{-5}$ | $5.83 \cdot 10^{-1}$ | $8.00 \cdot 10^3$ | $5.26 \cdot 10^{-6}$ | $4.40 \cdot 10^3$ | $9.55 \cdot 10^{-6}$ | $2.24 \cdot 10^3$ | $1.88 \cdot 10^{-5}$ | $1.52 \cdot 10^3$ | $2.77 \cdot 10^{-5}$ | $1.08 \cdot 10^3$ | $3.89 \cdot 10^{-5}$ |
| $9.0 \cdot 10^{-5}$ | $6.43 \cdot 10^{-1}$ | $9.33 \cdot 10^3$ | $2.80 \cdot 10^{-6}$ | $5.13 \cdot 10^3$ | $5.10 \cdot 10^{-6}$ | $2.61 \cdot 10^3$ | $1.00 \cdot 10^{-5}$ | $1.77 \cdot 10^3$ | $1.48 \cdot 10^{-5}$ | $1.26 \cdot 10^3$ | $2.08 \cdot 10^{-5}$ |
| $1.1 \cdot 10^{-4}$ | $6.88 \cdot 10^{-1}$ | $1.07 \cdot 10^4$ | $1.68 \cdot 10^{-6}$ | $5.87 \cdot 10^3$ | $3.05 \cdot 10^{-6}$ | $2.99 \cdot 10^3$ | $5.99 \cdot 10^{-6}$ | $2.03 \cdot 10^3$ | $8.83 \cdot 10^{-6}$ | $1.44 \cdot 10^3$ | $1.24 \cdot 10^{-5}$ |
| $1.3 \cdot 10^{-4}$ | $7.22 \cdot 10^{-1}$ | $1.20 \cdot 10^4$ | $1.09 \cdot 10^{-6}$ | $6.60 \cdot 10^3$ | $1.98 \cdot 10^{-6}$ | $3.36 \cdot 10^3$ | $3.88 \cdot 10^{-6}$ | $2.28 \cdot 10^3$ | $5.72 \cdot 10^{-6}$ | $1.62 \cdot 10^3$ | $8.05 \cdot 10^{-6}$ |
| $1.5 \cdot 10^{-4}$ | $7.50 \cdot 10^{-1}$ | $1.33 \cdot 10^4$ | $7.45 \cdot 10^{-7}$ | $7.33 \cdot 10^3$ | $1.35 \cdot 10^{-6}$ | $3.73 \cdot 10^3$ | $2.66 \cdot 10^{-6}$ | $2.53 \cdot 10^3$ | $3.92 \cdot 10^{-6}$ | $1.80 \cdot 10^3$ | $5.52 \cdot 10^{-6}$ |

## Conclusions

The difficulty in *K_d_* measurement forced the use of *K_m_* as an “apparent” equilibrium constant that could be used to assess enzyme-substrate affinity. This view is being challenged but the mechanistic meaning of *K_m_* has never been appropriately addressed. The possibility to be certain about the concentration of substrate providing (*in vitro*) half maximum V makes *K_m_* a valuable parameter in enzymology. This paper provides a further message, which is that *K_m_* is the constant defining the steady state, which is the actual state occurring in *in vitro* enzymatic assays aimed at *K_m_* determination. Further to *K_m_* and further to the specific property of an enzyme, i.e. its *k_cat_*, the substrate-enzyme association rate constant (*k_1_*) appears as relevant to understand the elasticity of the catalytic step in an overall system and to establish the period of time required to shift metabolic states, for instance in glycolysis from a resting situation to an anaerobic apnea. If we consider the fist enzyme in a metabolic route the flux after an increase in substrate availability is dependent on *k_1_* and not on *K_m_*. The same consideration would serve for a “Michaelian” transporter such as concentrative glucose transporters, the increase in glucose availability would increase the flux according to the association rate to the transporter and to the association rate of the first metabolic enzyme, the one producing glucose-6-phosphate (hexokinase or glucokinase).

In summary, the higher the *k_1_* value for enzymes in a given metabolism the higher the flux variation and the quicker the response to any perturbation. *k_1_* determination is a challenge that may be overcome by using stopped flow [7] or BIAcore equipment (see [25] for review) that allow real-time measuring of the association two molecules. Alternatively, *k_1_* may be deduced from progress [S]/t curves at different substrate concentrations [26,27]. Note that in such assays neither the *k_cat_* not the concentration of enzyme change; the development of novel numerical methods constitutes an alternative for calculating association rates from a set of progress [S] versus time curves.

